# LncRNA MIR4435-2HG mediates cisplatin resistance in HCT116 cells by regulating Nrf2 and HO-1

**DOI:** 10.1101/768986

**Authors:** Ping Luo, ShuGui Wu, ChaoMing Zhou, Xia Yuan, HongMi Li, JinPing Chen, YunFei Tian, Yang Qiu, XiaoMing Zhong

## Abstract

**PURPOSE:** Cisplatin resistance is still a serious problem in clinic. However, the underlying mechanism remains unclear. In this study investigated the drug resistance of cisplatin by the cisplatin resistance cell line HCT116R.

**RESULTS:** In this study, we found that LncRNA MIR4435-2HG level dramatically increased in the cisplatin resistance cell line HCT116R. Knockdown of MIR4435-2HG in HCR116R significantly restored the sensitivity to cisplatin, inhibited cell proliferation and promoted cell apoptosis. Furthermore, Nrf2 and HO-1 mRNA level, as the critical molecular of the oxidative stress pathway, was inhibited by the siRNA targeting to MIR4435-2HG, displaying MIR4435-2HG-mediated cisplatin resistance through the Nrf2/HO-1 pathway.

**CONCLUSION:** Our findings demonstrated that LncRNA MIR4435-2HG as a main factor could drive the cisplatin resistance of HCT116.

## Introduction

Colon cancer is one of the most common malignant tumors in the world. At present, there are more than 1 million new cases of colon cancer each year, which seriously endangers human life and quality of life[1].Most patients with metastatic colorectal cancer are still incurable. The positive response ratio of combined chemotherapy for colon cancer is only 20-47%, and most patients have recurrence[2].Cisplatin is widely used to treat varieties of cancers[3]. However, the drug resistance frequently occurs after a period of administration with effective response, and its specific mechanism is still not very clear[4].It has been shown that long non-coding RNA (lncRNA)interacts with chromatin regulatory proteins, RNA-binding proteins, and small RNAs to form a functional complex, regulating multiple important life processes[5-7].To investigate the role of lncRNA MIR4435-2HG in cisplatin resistance, we disrupted the expression of MIR44535-2HG in cisplatin resistant cells HCT116R and these cells became sensitive to cisplatin. At the same time, Nrf2 and HO-1 mRNA level also decreased.

## Materials and methods

### Cell culture and reagents

The colon cancer cell line HCT-116 was purchased from the American Type Culture Collection (Manassas, VA, USA). This is the cisplatin-sensitive cell lineand the cisplatin-resistant HCT-116R cell line was established from the parental HCT-116 cells by selection against cisplatin in the laboratory. Both HCT-116 and HCT-116R cells were maintained as a monolayer inMcCoy’s 5A medium (Gibco, Life Technologies, USA)supplemented with 10% fetal bovine serum, penicillin (100 U/ml),and streptomycin (100 μg/ml) in a humidified atmosphere of 5%CO2 at 37°C. Scramble and MIR4435-2HG siRNA were purchased from Thermo Fisher(China) and siRNA transfections were performed with Lipofectamine 2000 (LifeTechnologies).

### Cell proliferation assays

Cell proliferation was assessed with the WST-8 assay using Cell Counting Kit-8 (CCK-8) (DojindoLaboratories, Kumamoto, Japan). HCT-116 cells (5×10^3^/well) were seeded in 96-well plates and incubated for 24h. Cells wereincubated with or without 25μM cisplatin for 24h in McCoy’s 5A medium. Each treatment was carried out in triplicate. After incubation, 10 μl CCK-8 reagent was added to each well, and theplates were incubated at 37°C in an atmosphere of 5% CO2 for 4h. The optical density was then measured with a microplate reader(Multiskan JX; MTX Lab Systems, Vienna, VA, USA) at awavelength of 450 nm.

### Total RNA Isolation and Quantitative Real-Time Polymerase Chain Reaction (qRT-PCR)

Total RNA was isolated from treated cells usingthe TrizolReagent (Invitrogen, Carlsbad, CA),according to the manufacturer’s instruction. RNAconcentration and quality were determined by aspectrophotometer (Beckman, Brea, CA) and gelelectrophoresis, respectively. First-strand cDNA wassynthesized using the M-MLV Reverse Transcriptasekit (Invitrogen, Carlsbad, CA) in a total volume of20 mL. qRT-PCR analysis for gene expressionwas performed in triplicate using the SYBR GreenThe primer sequences used were MIR4435-2HGforward:

5′-CGGAGCATGGAACTCGACAG-3′,MIR4435-2HGreverse:

5′-CAAGTCTCACACATCCGGGC-3;Nrf2Forward:

5′-TACTCCCAGGTTGCCCACA-3′, Nrf2Reverse:

5′-CATCTACAAACGGGAATGTCTGC-3′;HO-1Forward:

5′-CACGCATATACCCGCTACCT-3′, HO-1Reverse:

5′-AAGGCGGTCTTAGCCTCTTC-3′; GAPDHForward:

5′-TATGATGATATCAAGAGGGTAGT-3′, GAPDHReverse:

5′-TGTATCCAAACTCATTGTCATAC-3′.

Caspase-3 activity assay. Cells were seeded into the 96 well plate at the density of 5000 and allowed to grow for 24 h. Then the siRNAs targeting to MIR4435-2HGwere transfected byLipofectamine RNAiMAX (ThermoFisher, China) according to the instruction manual. Cells were collected after 24 h and the caspase activity was determined using the Caspase-3 Assay Kit (Colorimetric) (Abcam ab39401, Cambridge, UK).

### Statistical analysis

Data are expressed as the mean±standard errorof the mean. Statistical analyses were performed using Prism Graphpad 6.0 (GraphPad Software, San Diego, CA, USA). Statistical significance was determined at p<0.05.

## Results

### The MIR4435 level is higher in HCT-116 cell line resistant to Cisplatin

First, we induced the drug resistance of HCT116 to Cisplatin by adding 25μm Cisplatin to HCT116 for 3 months, and we named it HCT-116R. The HCT-116R could proliferate with the presence of 25μm Cisplatin, while most HCT116 cells progressively die (Figure1A).We detected the response to cisplatin. Then RNA-Seq was used to analyze the changing cellular transcriptome of HCT116 and HCT116R. We found that the lncRNA MIR4435-2HG level dramatically increased about ten times and the result was verified with real-time PCR (Figure1B).

**Fig1.**
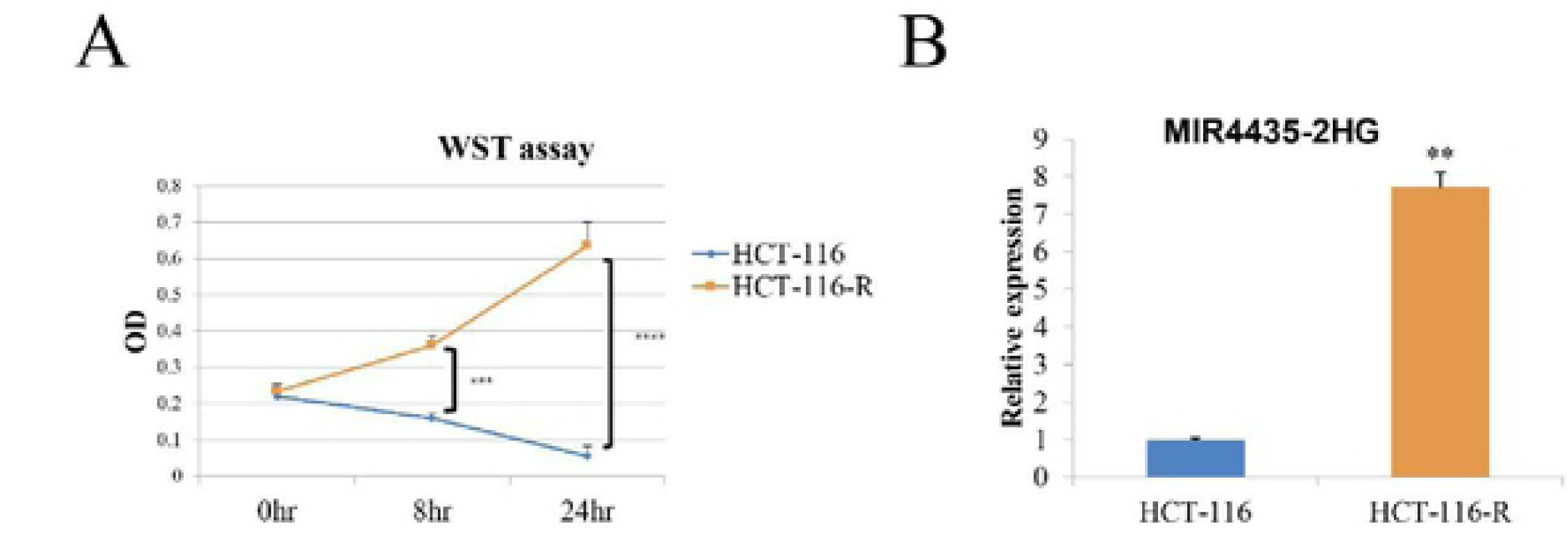
The MIR4435 level is higher in HCT-116 cell line resistant to Cisplatin. A. HCT116 and HCT116R cells were treated with 25µm Cisplatin, and the cell viability was tested by WST assay at 0, 8 and 24 hours later. B. The lncRNA MIR4435-2HG levels in HCT116 and HCT116R were detected by real time PCR. ** *P* < *0.01*, *** *P* < *0.001*, **** *P* < *0.0001*.

### Knockdown of MIR4435-2HG increased the sensitivity of HCT-116R to Cisplatin

To investigate the relationship of lncRNA MIR4435-2HG and cisplatin resistance, we knock down the lncRNA MIR4435-2HG by siRNA HCT116R by siRNA, and the MIR4435-2HG level dramatically decreased by siRNA, which were verified by real-time PCR (Figure2A).The knockdown of MIR4435-HG2 by siRNAincreased the sensitivity of HCT-116R to Cisplatin and the cell viability was tested by WST assay. With the presence of 25μm Cisplatin, the proliferation of HCT116R was inhibited by the siRNA targeting to the MIR4435-2HG, while there is no significant effect with the scramble sequence of siRNA (Figure2B). After knockdown of MIR4435-2HG, cisplatin increased casepase-3 activity about 2 times(Figure2C), and this indicated that the apoptosis of HCT116R.

**Fig2.**
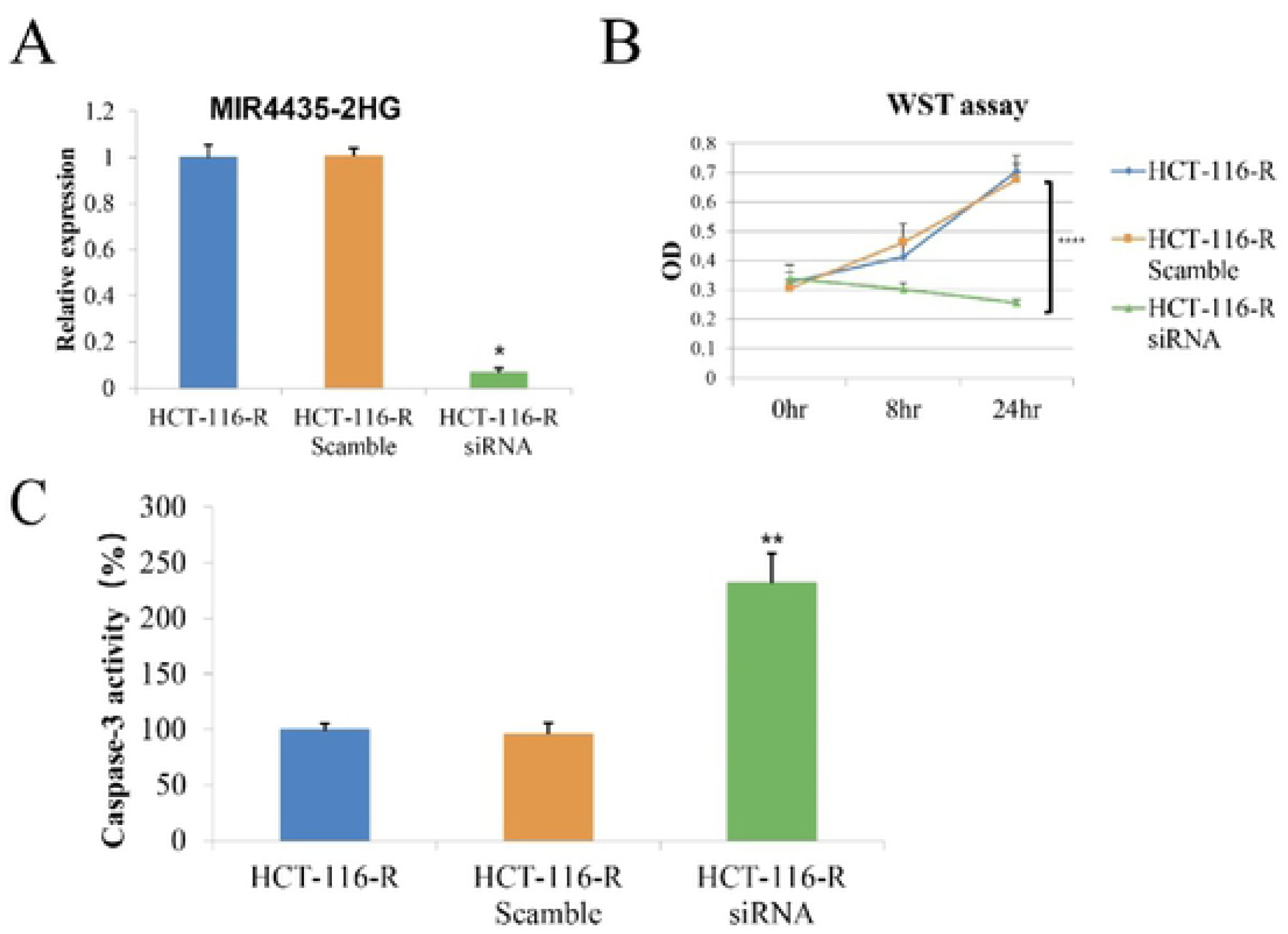
Transient knockdown of MIR4435 by siRNA sensitizes HCT-116Rto Cisplatin. A. The lncRNA MIR4435-2HG was knocked down in HCT-116R by siRNA, and the effect of siRNA were verified by real-time PCR. 8. The knockdown of MIR4435-HG2 by siRNAincreased sensitivity of HCT-116R to Cisplatin and the cell viability was tested by WST assay at 0, 8 and 24 hours after the administration of Cispalin. C The casepase-3 activity was tested after the Cisplatin treatment 24 hours later. The data are presented as means ± SDs. * *P* < *0.05*, ** *P* < *0.01*, **** *P* < *0.0001*.

### Knockdown of MIR4435-2HG decreased the Nrf2 and HO-1 mRNA level related to oxidative damage

Nuclear factor erythroid 2-related factor 2 (Nrf2) is a keytranscription factor that regulates antioxidant anddetoxification enzymes and Heme oxygenase-1 (HO-1) is a Nrf2-regulated gene that plays a critical role in the prevention of inflammation. Transfection of MIR4435-2HG siRNA downregulated the mRNA levels of Nrf2 and HO1 significantly after the 25μM cisplatin treatment (Figure3), and this phenomenon indicated the role of MIR4435-2HG is involved the oxidative stress.

**Fig3.**
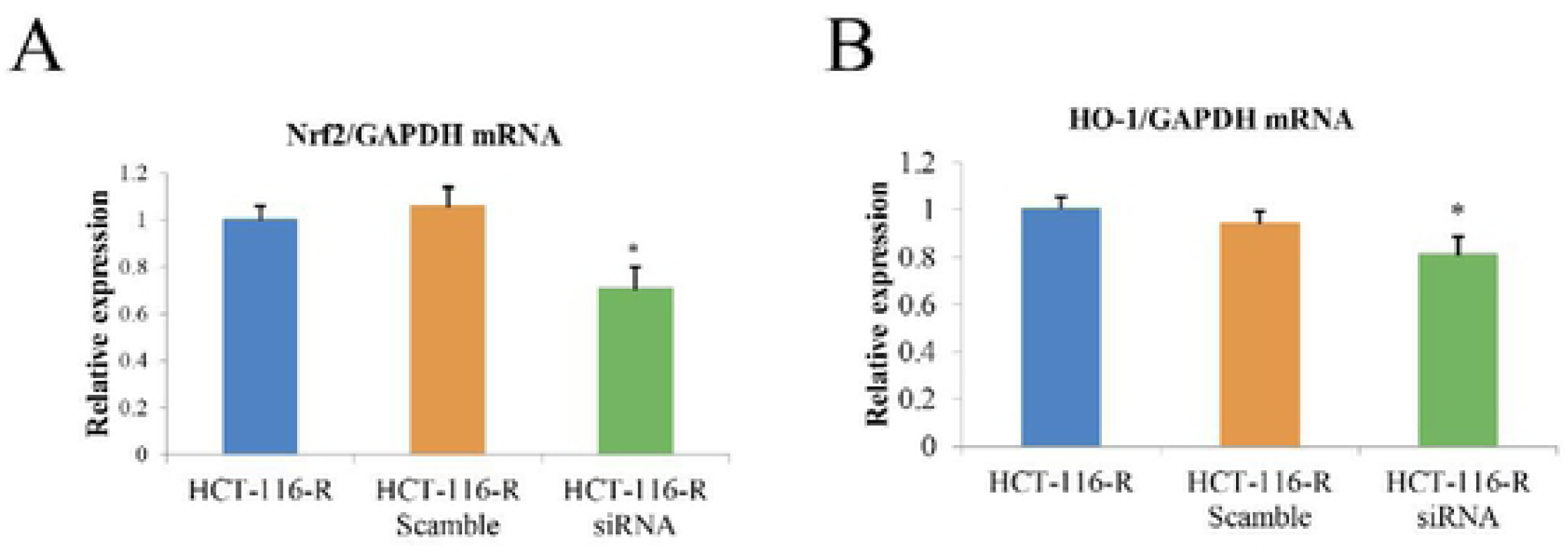
Knockdown of MIR4435 decreased the NRF2 and HO-1 mRNA level related to oxidative damage. The relative NRF2 (A) and HO-1 mRNA (B) levels of HCT-116R cells were analyzed by real time PCR and the results were normalized to the reference geneGAPDH. The data are presented as means ± SDs. * *P* < *0.05*

## Discussion

Cisplatin is thought to formulate the covalent adducts with some bases in the DNA, then could cure many testicular and some ovarian carcinomas[8, 9]. However, the efficiency of cisplatin is low in colorectal cancer(CRC), with fewer than 20% clinical responses whenused alone or in combination with other anti-cancer drugs[10]. Last few years, several studies suggested that some lncRNA contributed the cisplatin resistance via various different mechanism[11].

LncRNA MIR4435-2HG has been reported to related with the hepatocellular carcinoma[12] and lung cancer[13]. Wen et al. found the lncRNA-MIR4435-2HG might participate in the development of colorectal cancer via the P38/MAPK and VEGF pathway[14].In this report, we found that lncRNA MIR4435-2HG increased about 7-8 folds in cisplatin resistant cell line compared to normal cell line, which implied that MIR4435-2HG may involve the cisplatin resistant. Then we knocked down the MIR4435-2HG level with siRNA, and about 90% of MIR4435-2HG expression was inhibited by siRNA in cisplatin resistant HCT-116R cells. The effect of the MIR4435-2HG knock down in HCT-116R cells is that 24 hours cisplatin administration caused the about one third of HCT-116R death, as a control the HCT-116R cells without the MIR4435-2HG expression disruption proliferate about two folds in 24hours. The cisplatin resistant HCT-116R cells became sensitive to cisplatin after the MIR4435-2HG knock down. Besides that, after the 24h cisplatin treatment, compared to the HCT-116R cells, the caspase-3 activity increased 2.5 folds in the MIR4435-2HG knockdown cells, while the scramble siRNA did not have any effect. It indicated that the cisplatin induced cell death may be mainly produced by the apoptosis.

Nuclear factor erythroid 2-related factor 2 (Nrf2), a transcription factor, responses for the oxidative stresses and plays a key role in redox homeostasis[15]. Heme oxygenase (HO), acting as downstream of Nrf2[16], is an enzyme that catalyzes the degradation of heme[17, 18]. These two components usually increased in different types of tumors and correlate with tumor progression, aggressiveness, resistance to therapy, and poor prognosis[19]. In our system, the knockdown of MIR4435-2HG decreased the NRF2 and HO-1 mRNA level significantly.

## Conclusions and Perspectives

This report we observed that lncRNA MIR4435-2HG as a main factor could drive the cisplatin resistance of HCT116.Moreover, lncRNA MIR4435-2HG might participate in the development of cisplatin resistance through the Nrf2/HO-1 pathway. The current study unveiled the roles of lncRNA MIR4435-2HG in colorectal cancer cisplatin resistance and provided the underlying mechanism of cisplatin resistance. Based on the mechanisms underlying cisplatin resistance, the following strategies have been proposed to overcome the resistance in colorectal cancer in the future: 1) develop new platinum drugs;2) improve cisplatin delivery to tumor; 3) specifically target cisplatin resistance mechanisms; and 4) combine cisplatin with other drugs. Understanding the molecular mechanisms underlying cisplatin resistance may be helpful to identify patients with better treatment, and thus oncologists may be able to provide an effective therapy for those patients.

## Acknowledgements

Not applicable.

## Abbreviations 缩略语

lncRNAs: Long non-coding RNAs
Nrf2: Nuclear factor erythroid 2-related factor
HO-1: Heme oxygenase-1
qRT-PCR: Quantitative Real-Time Polymerase Chain Reaction
CRC: colorectal cancer

## Authors’ contributions

XMZ and YQ conceived and designed the experiments, reviewed drafts of the paper. Y Q and PL participates in Statistical analysis, analyzed and interpreted the patient data. SGW and CMZ carried out the study, collected and analyzed the data. X Y and HM Lwas a major contributor in writing the manuscript. JPC is mainly responsible for making figure, and YFT contributed reagents/materials/analysis tools. All authors read and approved the final manuscript.

## Availability of data and materials

The analyzed datasets generated during the study are available from the corresponding author on reasonable request.

## Ethics approval and consent to participate

The present study was approved by the Ethics Committee of Jiangxi province Cancer Hospital.

## Consent for publication

Not applicable.

## Footnotes

Ping Luo, ShuGui Wu and ChaoMing Zhou contributed equally to this work.

## Contributor Information

1.Ping Luo.

2.ShuGui Wu.

3.ChaoMing Zhou.

4.XiaYuan.

5.HongMi Li.

6.JinPing Chen.

7.YunFei Tian.

8.Yang Qiu.

9.XiaoMing Zhong.

## Reference

1. Benson AB 3rd, Venook AP, Cederquist L, Chan E, Chen YJ, Cooper HS, Deming D, et al. Colon cancer, version 1.2017, NCCN clinical practice guidelines in oncology. Journal of the National Comprehensive Cancer Network, 2017. 15: 370–398.

2. Grothey A, Van Cutsem E, Sobrero A, Siena S, Falcone A, Ychou M, Humblet Yet al, et al. Regorafenib monotherapy for previously treated metastatic colorectal cancer (CORRECT): an international, multicentre, randomised, placebo-controlled, phase 3 trial. Lancet, 2013. 381: 303–312.

3. Amable L. Cisplatin resistance and opportunities for precision medicine. Pharmacol Res, 2016. 106: 27–36.

4. Galluzzi L, Vitale I, Michels J, Brenner C, Szabadkai G, Harel-Bellan A, Castedo M, et al. Systems biology of cisplatin resistance: past, present and future. Cell Death & Disease, 2014: e1257.

5. Quinodoz S, M Guttman. Long noncoding RNAs: an emerging link between gene regulation and nuclear organization. Trends Cell Biol, 2014. 24: 651–63.

6. Zhu, J,Fu H, Wu Y, Zheng X.Function of lncRNAs and approaches to lncRNA-protein interactions. Science China Life Sciences, 2013. 56:876–885.

7. Long Y,Wang X, Youmans DT, Cech TR. How do lncRNAs regulate transcription? Sci Adv, 2017. 3: eaao2110.

8. Galluzzi L,Senovilla L, Vitale I, Michels J, Martins I, Kepp O, Castedo M, et al. Molecular mechanisms of cisplatin resistance. Oncogene, 2012. 31: 1869–83.

9. Makovec T. Cisplatin and beyond: molecular mechanisms of action and drug resistance development in cancer chemotherapy. Radiol Oncol, 2019. 53: 148–158.

10. Hammond WA, A Swaika, K Mody. Pharmacologic resistance in colorectal cancer: a review. Ther Adv Med Oncol, 2016.8: 57–84.

11. Hu Y, Zhu QN, Deng JL, Li ZX, Wang G, Zhu YS,et al., Emerging role of long non-coding RNAs in cisplatin resistance. Onco Targets Ther, 2018. 11: 3185–3194.

12. Kong Q, Liang C, Jin Y, Pan Y, Tong D, Kong Q, Zhou J. The lncRNA MIR4435-2HG is upregulated in hepatocellular carcinoma and promotes cancer cell proliferation by upregulating miRNA-487a. Cell Mol Biol Lett, 2019. 24: 26.

13. Qian H, Chen L, Huang J, Wang X, Ma S, Cui F, Luo L,et al.The lncRNA MIR4435-2HG promotes lung cancer progression by activating beta-catenin signalling. J Mol Med (Berl), 2018. 96: 753–764.

14. Ouyang W, Ren L, Liu G, Chi X, Wei H.LncRNA MIR4435-2HG predicts poor prognosis in patients with colorectal cancer. PeerJ, 2019. 7: e6683.

15. Truyen Nguyen, Paul Nioi, Cecil B. Pickett.The Nrf2-antioxidant response element signaling pathway and its activation by oxidative stress. J Biol Chem, 2009. 284:13291–5.

16. Na HK, Surh YJ. Oncogenic potential of Nrf2 and its principal target protein heme oxygenase-1. Free Radic Biol Med, 2014. 67:353–65.

17. Tenhunen R, HS Marver,R Schmid. The enzymatic conversion of heme to bilirubin by microsomal heme oxygenase. Proc Natl Acad Sci U S A, 1968. 61: 748–755.

18. Loboda A, Damulewicz M, Pyza E, Jozkowicz A, Dulak J,et al. Role of Nrf2/HO-1 system in development, oxidative stress response and diseases: an evolutionarily conserved mechanism. Cell Mol Life Sci, 2016. 73: 3221–47.

19. Furfaro AL, Traverso N, Domenicotti C, Piras S, Moretta L, Marinari UM, Pronzato MA,et al.The Nrf2/HO-1 Axis in Cancer Cell Growth and Chemoresistance. Oxid Med Cell Longev, 2016. 2016:1958174.

